# PatchMAN docking: Modeling peptide-protein interactions in the context of the receptor surface

**DOI:** 10.1101/2021.09.02.458699

**Authors:** Alisa Khramushin, Tomer Tsaban, Julia Varga, Orly Avraham, Ora Schueler-Furman

## Abstract

Peptide docking can be perceived as a subproblem of protein-protein docking. However, due to the short length and flexible nature of peptides, many do not adopt one defined conformation prior to binding. Therefore, to tackle a peptide docking problem, not only the relative orientation between the two partners, but also the bound conformation of the peptide needs to be modeled. Traditional peptide-centered approaches use information about the peptide sequence to generate a representative conformer ensemble, which can then be rigid body docked to the receptor. Alternatively, one may look at this problem from the viewpoint of the receptor, namely that the protein surface defines the peptide bound conformation.We present PatchMAN (Patch-Motif AligNments), a novel peptide docking approach which uses structural motifs to map the receptor surface with backbone scaffolds extracted from protein structures. On a non-redundant set of protein-peptide complexes, starting from free receptor structures, PatchMAN successfully models and identifies near-native peptide-protein complexes in 62% / 81% within 2.5Å / 5Å RMSD, with corresponding sampling in 81% / 100% of the cases, outperforming other approaches. PatchMAN leverages the observation that structural units of peptides with their binding pocket can be found not only within interfaces, but also within monomers. We show that the conformation of the bound peptide is sampled based on the structural context of the receptor only, without taking into account any sequence information. Beyond peptide docking, this approach opens exciting new avenues to study principles of peptide-protein association, and to the design of new peptide binders.

## Introduction

Peptide-protein interactions - namely interactions mediated by short segments or motifs often located in disordered regions - are very common in the cell, constituting about 40% of the overall protein interactions (1). Such interactions participate in many important cellular processes like regulation and cell-signaling (2). Therefore structural characterization of such interactions is crucial for the understanding of many biological pathways and their potential in the development of therapeutic targets and other biotechnological applications (3). However, such interactions are often weaker and more transient than globular protein interactions and therefore more challenging to characterize experimentally, highlighting the need for developing computational tools for modeling their structures.

The intuitive way to look at *protein-peptide* docking is as a sub-problem of *protein-protein* docking. However, this approach presents several hurdles, since in addition to the problem of finding the relative orientation between the two partners, the peptide conformation is often not known or does not even assume a defined structure before binding the receptor (4). When the binding site is known and a coarse model of a peptide-protein complex is available, it can be further refined to high accuracy by local refinement protocols, such as Rosetta FlexPepDock developed by us (5, 6). In the absence of such information however, global docking has to be performed. To reduce the conformational space needed to sampling both the peptide conformation and its location on the receptor, many currently existing peptide-docking approaches tackle this problem by decoupling the folding and docking steps, generating a peptide conformational ensemble for subsequent docking (7–10). For example in the PIPER-FlexPepDock (PFPD) protocol previously developed by us (11), a conformer ensemble is generated using the Rosetta Fragment Picker (12) (similar to the first step in traditional *ab initio* folding (13)). This ensemble is then rigid body docked using PIPER (14) and further refined by FlexPepDock. This approach is also implemented in the InterPep docking protocol (15). MDockPeP uses sequence-similar fragments extracted from monomers (16), while in HADDOCK and pepATTRACT, peptide conformations are represented by idealized secondary structure fragments (8, 10), and the CABS-dock protocol uses random peptide conformations for subsequent docking and refinement (17). All these approaches are united by the idea that the peptide, as a separate protein, carries enough information for its separate folding, or at least the determination of a conformer ensemble that represents its conformational preferences. But what if the conformational ensemble of the peptide does not include the conformation that it adopts upon binding? In such a case, the rigid-body step of the docking protocol will not be able to fit the peptide into the binding pocket. An alternative solution for finding the bound conformation of the peptide is template-based modeling. Many protein-protein interactions can be modeled based on a solved structure of a homolog complex (18), and the same can be applied to protein-peptide interactions (19). However, such an approach is restricted to a limited amount of solved protein-peptide complexes.

We present here a novel approach for blind peptide docking, which we name PatchMAN (Patch-Motif AligNments), that combines a global search with template-based modeling, benefitting from both strategies. We look at peptide docking from the viewpoint of the *receptor*, building on the assumption that the protein surface carries enough information to determine the peptide bound conformation. This is based on the previously proposed theory that peptide-protein interactions often mimic structural characteristics that are typical to monomeric folds (20), hinting at a large reservoir of information that can further be used for peptide-protein docking. PatchMAN uses surface patches, defined as bundles of disjoint backbone segments, to search for similar “pockets” that contain a peptide stretch interacting with it in a dataset of protein structures that includes monomers as well as protein-protein and protein-peptide complexes. The backbone conformation of such peptide stretches is then superimposed back to the receptor protein, and is used as a starting point for local peptide docking refinement.

PatchMAN shows performance superior to current peptide-docking methods, including our recent implementation of AlphaFold2 (21) for peptide docking (22). As such, PatchMAN opens new opportunities to model more complicated protein-peptide-like interactions, in addition to facilitating design of new peptide binders.

## Results

### General overview of the PatchMAN approach: docking by globally mapping the receptor surface with local motif templates

In general, protein-peptide as well as protein-protein interaction modeling can be split into two categories: template-based modeling, in which new interactions are modeled based on solved structures of similar interactions, and free modeling, in which a large number of new rigid body orientations and internal degrees of freedom are sampled. In PatchMAN we suggest combining the two, by generating peptide templates on the whole protein surface, thus sampling the binding sites and “folding” the peptide at the same time. The protocol consists of 4 consecutive steps (**Figure 1**): (1) Definition of surface patches on the receptor; (2) Identification of structural motif matches in protein structures, and an interacting fragment that can be used as template for the bound peptide; (3) Generation of the peptide-protein complex template structure, by superimposing the interacting peptide back onto the receptor, and (4) Replacing side chains according to the peptide sequence (threading), refinement and scoring of the model.

**Figure 1.**
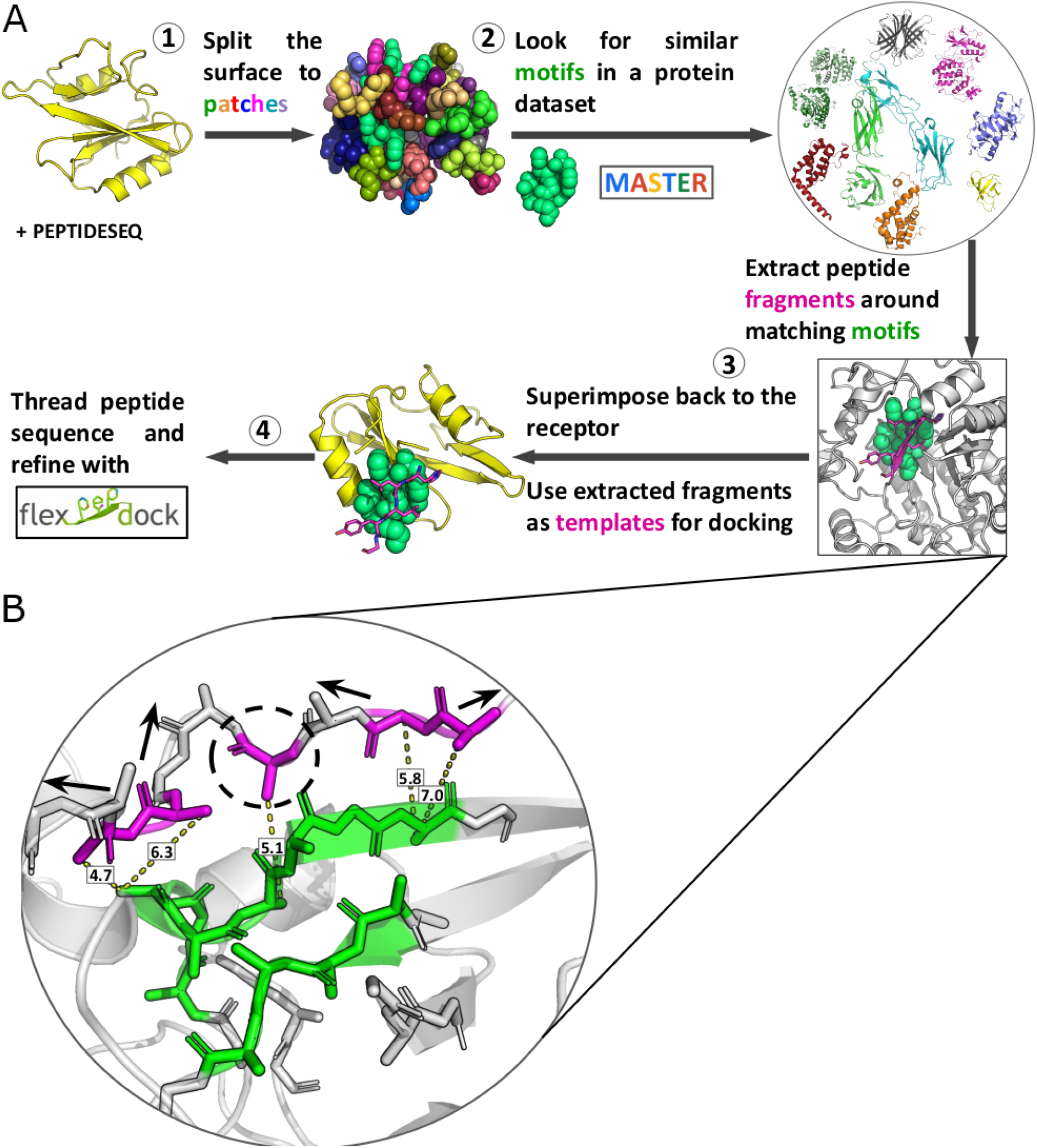
The PatchMan protocol. **(A) Flowchart.** The input is a receptor PDB file and a peptide sequence. (1) *Definition of surface motifs on the receptor:* The protein surface is defined based on solvent accessibility, and then split into small structural surface patches. (2) *Identification of structural matches in protein structures:* Matches are detected using MASTER search against a non-redundant dataset of protein structures; (3) *Generation of the peptide-protein complex structure:* the peptide fragment is determined (see (B)) and superimposed onto the receptor. Then the peptide sequence is threaded onto the identified complementing fragment; (4) *Refinement and scoring:* the initial structures are refined using the Rosetta FlexPepDock refinement protocol, and top-scoring models are selected as final predictions. **(B) Extracting peptide fragments.** Neighboring residues (magenta) around the matching motif (green) are defined with Cβ distance within 8Å of the motif. Consecutive backbone stretches are then elongated in both directions to the desired peptide length. Arrows indicate stretches that can be elongated. Single residue (indicated with dashed circle) will not be elongated. See Methods for more details.

In the following we describe the protocol in more details (see also Methods section). For the sampling step, we first identify the surface residues based on surface accessible area (SASA). Next, the surface is split into patches consisting of one or more peptide segments. Those patches are then used to search for similar motifs in a diverse non-redundant database of protein structures (maximum 30% pairwise sequence identity), using MASTER (23). Peptide stretches around every found motif are extracted (see **Figure 1B**). If an interacting fragment is shorter than the required peptide length, it is elongated in both directions, so that even patches only partially covering the binding site can lead to generation of a near-native template. The extracted peptide fragments are then superimposed back to the receptor protein using the rotational matrices from the patch-motif alignment. At this point the receptor protein surface is fully mapped with templates for local peptide docking. Once the sampling is complete, the peptide sequence is threaded onto the generated peptide templates. These starting structures are then refined using Rosetta FlexPepDock (5). Finally, all models are scored and the best models are selected. Additional and more extensive details on each step, including specific parameters, are described in the Methods section.

### PatchMAN performance

For the initial estimation of the method performance we ran PatchMAN on a non-redundant dataset containing 26 solved protein-peptide complexes previously used to assess performance of PIPER-FlexPepDock (PFPD) (11). It includes two subsets of complexes: one with known binding motifs (here the motifs are ELMs - Eukaryotic Linear Motifs (24)) and the second for which no motifs have (yet) been reported. For all the complexes, free (unbound) receptor structures are available, and those were used in the current study to reflect a blind, real-world scenario. The structures of the solved complex were filtered out from the template set.

PatchMAN generates and identifies for 81% / 62% of the complexes within the top 10 cluster representatives a near-native model within 5 Å / 2.5 Å RMSD, respectively (see **Table I, Figure 2A-C**, and **Supplementary Figure S1**). It outperforms other methods, including PFPD, our application of AlphaFold2 to peptide docking (22), InterPep(15) MDockPeP(16), HADDOCK (8, 10), pepATTRACT (8, 10), and CABS-dock (17), which are based on a variety of approaches (as detailed in the Introduction) (**Figure 2A**).

**Figure 2:**
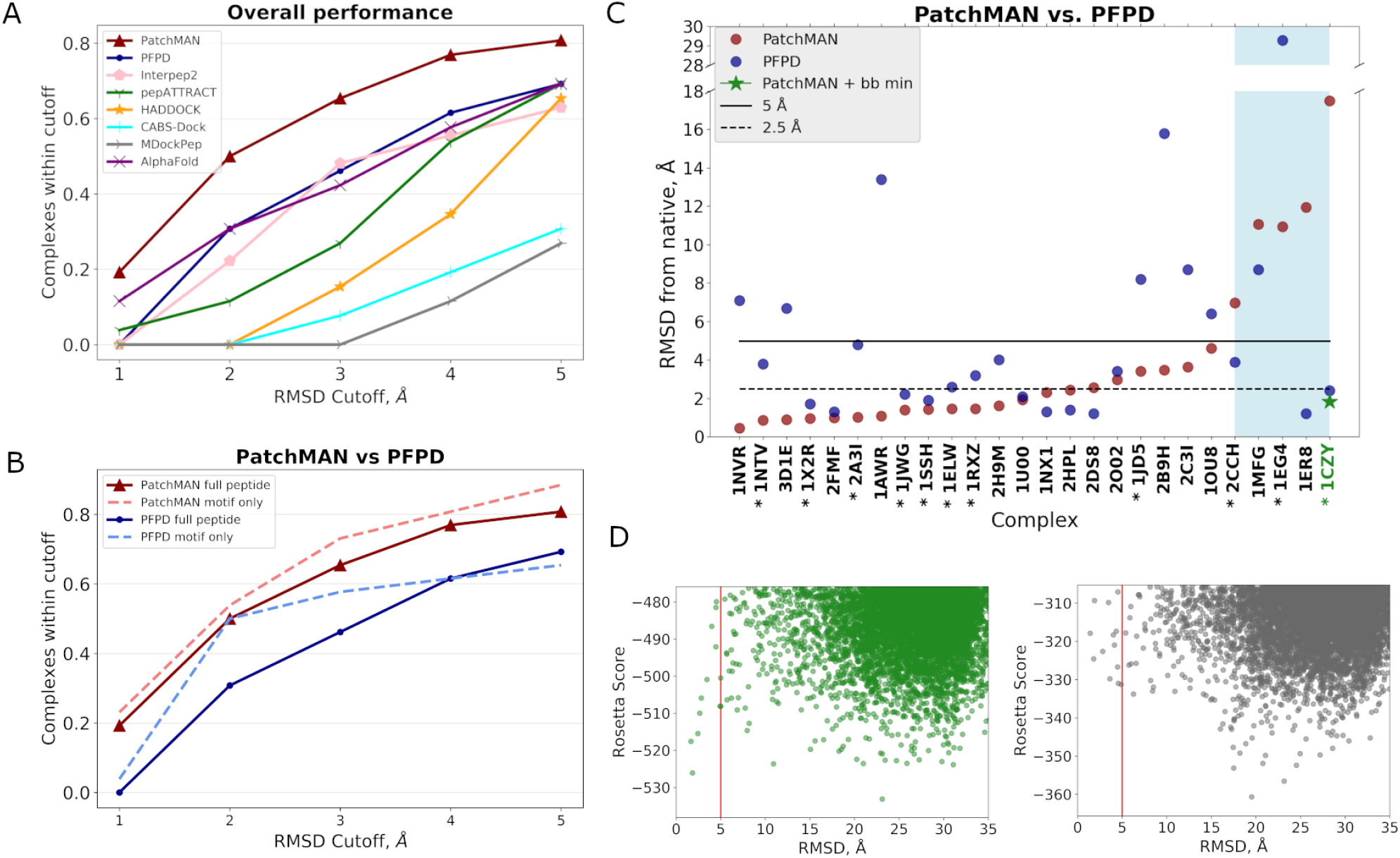
Highly accurate modeling of peptide-protein complexes with PatchMan. **A**. Comparison of PatchMAN to other approaches. For each method the y-axis shows the cumulative success, namely the fraction of complexes modeled within the RMSD threshold indicated in the x-axis. The top-performing model is considered for each complex (*i.e.*, the best RMSD among top 10 cluster representatives). PatchMAN performance is superior to all other methods on this dataset (see text for more details). **B.** Modeling only the motif sequence (dashed lines; extracted from the full peptide sequence) significantly improves performance of PIPER FlexPepDock (PFPD) but only slightly affects PatchMAN performance. **C.** Detailed comparison of PatchMAN and PFPD performance. PatchMAN RMSD values are plotted in red, PFPD in blue. Shaded region of the plot indicates complexes for which PatchMAN failed to produce models within 5Å RMSD, as for example the 1CZY complex (25), highlighted in green and described in (D). **D.** Including receptor flexibility in the refinement step can resolve failed docking, as shown for 1CZY: Near-native conformations (left to the highlighted red line) are only identified in a simulation with receptor minimization (green, right), but not in the corresponding refinement that keeps the receptor backbone fixed (grey, left). In all plots the RMSD measure reported is backbone interface RMSD.

**Table I:** summary of performance for a representative, non-redundant benchmark (from (11)).

| Complex PDB ID | Unbound receptor | Best model* | Best sampled RMSD | Peptide length |
| --- | --- | --- | --- | --- |
| 1AWR | 2ALF | 1.1 | 0.7 | 6 |
| 1ER8 | 4APE | 11.9 | 3.1 | 8 |
| 1NVR | 2QHN | 0.4 | 0.4 | 5 |
| 1NX1 | 1ALV | 2.3 | 1.0 | 11 |
| 1OU8 | 1OU9 | 4.6 | 2.7 | 8 |
| 1U00 | 2V7Y | 1.9 | 1.9 | 9 |
| 2B9H | 2B9F | 3.5 | 2.2 | 12 |
| 2C3I | 2J2I | 3.6 | 1.7 | 8 |
| 2DS8 | 2DS7 | 2.6 | 1.2 | 6 |
| 2FMF | 1JBE | 1.0 | 1.0 | 13 |
| 2H9M | 2H14 | 1.6 | 1.3 | 5 |
| 2HPL | 2HPJ | 2.4 | 2.1 | 5 |
| 2O02 | 2BQ0 | 3.0 | 3.0 | 14 |
| 3D1E | 3D1G | 0.9 | 0.9 | 6 |
| 1CZY | 1CA4 | 17.5 | 1.3 | 7 |
| 1EG4 | 1EG3 | 10.9 | 3.8 | 13 |
| 1ELW | 1A17 | 1.4 | 1.1 | 8 |
| 1JD5 | 1JD4 | 3.4 | 2.3 | 8 |
| 1JWG | 1JWF | 1.4 | 1.2 | 5 |
| 1MFG | 2H3L | 11.1 | 1.9 | 9 |
| 1NTV | 1P3R | 0.9 | 0.9 | 10 |
| 1RXZ | 1RWZ | 1.5 | 1.1 | 11 |
| 1SSH | 1OOT | 1.4 | 0.9 | 11 |
| 1X2R | 1X2J | 1.0 | 0.6 | 9 |
| 2A3I | 2AA2 | 1.0 | 1.0 | 12 |
| 2CCH | 1H1R | 7.0 | 3.4 | 12 |

\* top-RMSD model among 10 top-scoring clusters

A more detailed comparison to the PFPD blind peptide docking protocol, that also uses FlexPepDock in the refinement step, reveals that while PFPD performs relatively poorly for full-length peptides, and much better for the motif only docking simulations, PatchMAN performs similarly well both when modeling full-length peptides containing flanking regions and when only the peptide motifs are used (**Figure 2B**). In most cases PatchMAN outperforms PFPD (**Figure 2C**), except for 2 cases - 1CZY and 1ER8. We checked what caused PatchMAN to fail on these two examples. We noticed that for 1CZY PatchMAN sampled near-native complexes, but failed to identify them in the scoring step (**Figure 2D** left panel). We show that adding receptor backbone minimization to the refinement step solves this issue immediately (**Figure 2D** right panel), as it does for the PFPD runs (of note, all PFPD results shown here included receptor backbone minimization in the refinement step). For 1ER8 however, we observed no sampling at the binding site (see **Supplementary Figure S1**). We found that this is due to the absence of matching motifs for the patches covering the binding site. This can be solved by either further fine-tuning of the patch definition, increasing the template dataset, or loosening the RMSD threshold for the match search.

### The receptor surface can be mapped by local structural motif matches

Our results demonstrate that even with a relatively small non-redundant set of proteins, we can model the conformation of the peptide bound to its receptor (**Figure 2**). We show that in all cases PatchMAN samples peptide conformations within 5Å RMSD from the native complex structure. This is one of the many possible conformations generated that cover the whole receptor surface (See **Supplementary Figure S2**, and the energy landscapes presented throughout this paper and in **Supplementary Figure S1**). This implies that even within a limited dataset of proteins there are many motifs which are similar to receptor surface patches, and which include complementing peptide stretches fitting into these surface pockets. Increasing the size of the database for the template search can help to introduce more diversification of the sampling. More diverse motifs will help finding less trivial matches, thus introducing more intrinsic flexibility to the receptor and aiding in solving more complicated cases.

We analyzed the sequence similarity between the peptide templates and the docked peptide that led to generation of the near-native models (top 1% lowest scoring models within 5Å RMSD; **Figure 3A**). We found that sequence identity of the peptide templates is predominantly below 30%, with many templates showing no sequence identity to the native peptide. For most cases (15/21), the best model is derived from a template with less than 30% sequence identity. These results indicate that peptide bound conformation can be sampled based on the receptor surface conformation only, without regard to the peptide sequence.

**Figure 3.**
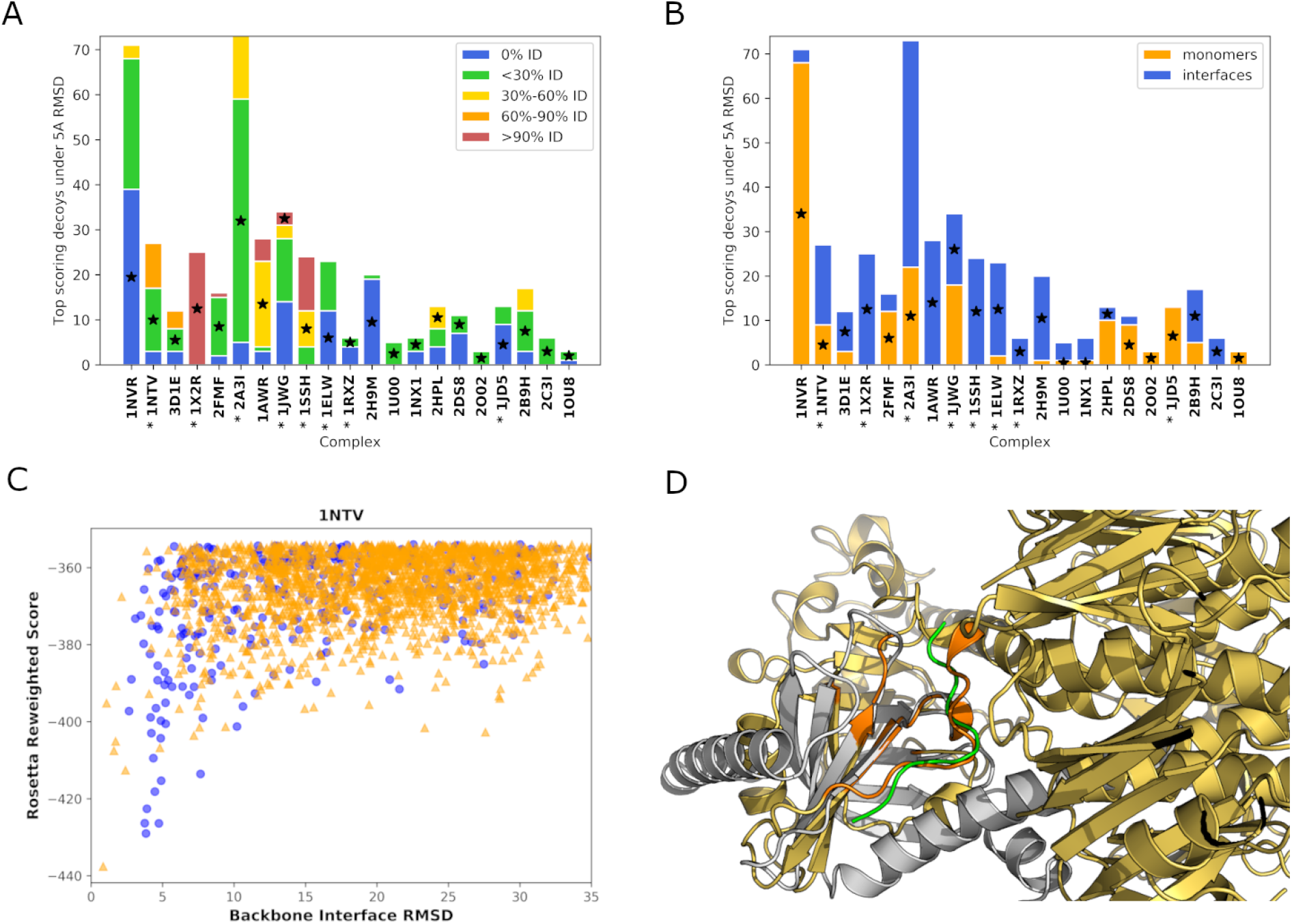
Peptide templates leading to high-resolution models are very varied: They show no sequence similarity and can be extracted from monomers. **A.-B.** Detailed results for different complexes. Shown are the top-scoring models (1%) within 5Å RMSD, with stars representing the best RMSD model (identified in the top 10 cluster representatives). Complexes are sorted in increasing order of the best RMSD. The structures from the “motif” set are marked with an asterix. **A.** Most of the top-scoring near-native models are modeled using templates with very low sequence identity. **B.** The source of low-RMSD templates comes from monomers (orange) as well as interfaces (blue). **C.-D.** Details of the prediction for 1NTV (Disabled-1 (Dab1) PTB domain-ApoER2 peptide complex) (26): (**C**) Energy landscape. Models generated based on templates originating from monomers and interfaces are indicated in orange triangles and blue circles, respectively. See **Supplementary Figure S1** for more energy landscapes. (**D**) Structure of the interaction, together with the template that was used for modeling (PDB ID 1LTI, Heat labile enterotoxin type I (27)). The free receptor structure (1P3B (28)) is shown in grey, the native peptide in green, the monomer from which the template was extracted in gold. The matching motif and peptide template are colored in orange.

### Many templates for near-native models can be extracted from monomer structures

Analysis of the templates revealed that although most of the templates originate from monomers (an average of 77%) there is a great diversity in the source of the templates that leads to the final near-native complex, depending on the type of interaction (See **Figure 3B** and **Supplementary Figure S1**). For half of the cases (10/21) the best model is derived from a monomer template structure. In many cases successful templates were extracted from both interfaces and monomers, where the interfaces include both peptide-protein complexes as well as protein-protein complexes. However, for several cases (1NTV, 1U00, 2FMF) the only near-native complexes originated from monomers. We will look more into detail of the PatchMAN prediction of the 1NTV complex (Disabled-1 (Dab1) PTB domain-ApoER2 peptide complex) (**Figure 3C,D** and **Figure 4**). The top-scoring structure (RMSD = 0.85Å; **Figure 3C**) was generated using a template extracted from the structure of the monomer of Heat labile enterotoxin type I (**Figure 3D**). This is an example of non-trivial template extraction that requires finding a “non-perfect match”: in this case the patch consists of 4 disjoint segments (patch-motif RMSD = 1.1Å). This is in contrast to cases where homologous complexes of a peptide-protein interaction are available as a template (as, e.g., for 1X2R). These results demonstrate that protein monomers can indeed serve as models for peptide conformations and should be utilized in peptide-protein docking.

**Figure 4.**
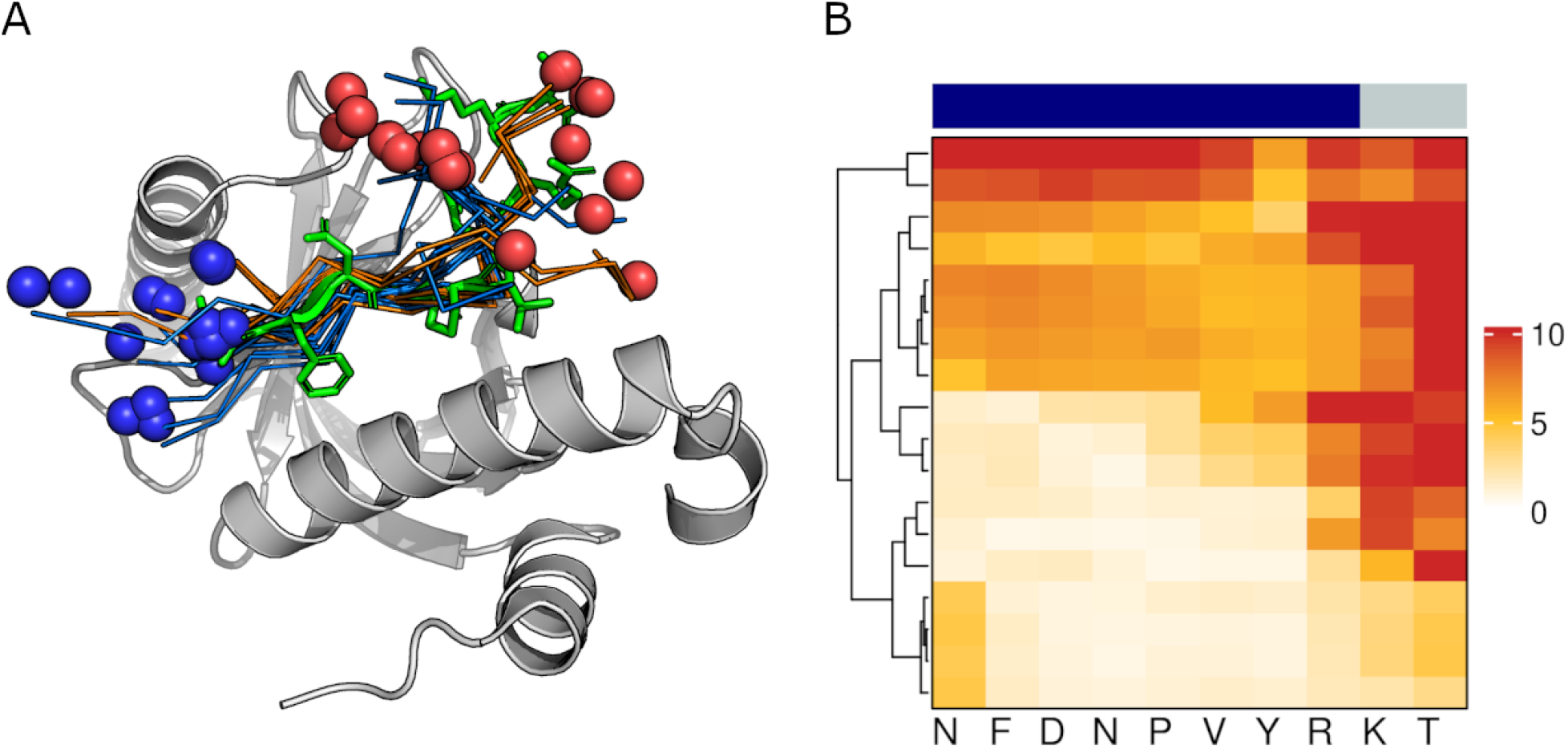
Sampling at the binding site of 1NTV shows local diversity. **A.**Templates extracted for a patch defining the binding site. In ribbon - templates, green sticks representation - native peptide, N and C termini in blue and red spheres accordingly. For each template the best stretch (out of a few sliding windows) is shown. Template coloring is the same as in **Figure 3A** depending on whether it comes from monomer/interface. **B.** Backbone per-residue RMSD for the templates shown in (A). The upper bar indicates the motif and the flanking region with blue and grey color, respectively.

Inspection of the extracted templates at the 1NTV binding site reveals that they show considerable variability (**Supplementary Figure 2C**). Zooming into the templates selected for the specific peptide sequence (*i.e.*, looking at the best stretch out of several sliding windows extracted from a protein including the matching motif) shows convergence towards the near-native peptide conformation, and does not include conformations of different secondary structure or opposite orientation (**Figure 4A**). At the same time we see a local diversity of the templates in the binding site with per-residue RMSD that ranges up to 5Å for the motif part (**Figure 4B**).

### PatchMAN overcomes conformational changes induced by ligand binding: The FERM domain example

One of the big challenges in protein docking and in peptide docking specifically is that the binding pocket can undergo conformational changes upon binding of the ligand. Given that at the sampling stage we use only the receptor surface information, it is crucial that the representation of the surface will be robust to such changes. This challenge is addressed in PatchMAN in two ways: (1) Backbone-based search: the surface patches that we use for screening of matching motifs are represented as bundles of backbone segments, thus allowing for flexibility at the side-chain level with surface rotamers. (2) Diversification of matches: For each surface patch we use matching motifs with very low RMSD for finding easy templates (e.g. homologous structures), but also more distant motifs with RMSD ~1.5Å to capture cases of possible backbone conformational changes.

One example of a protein that undergoes such conformational change is the Moesin FERM domain (**Figure 5A**). Prior to binding, the F3b binding pocket is closed and inaccessible to the peptide (29, 30). However, in case a binding site is known, the pocket could be opened by positioning peptide into the binding site, with subsequent refinement of the structure, as implemented previously e.g. in CAPRI target T121 (31). To test the ability of PatchMAN to deal with such conformational changes we attempted to dock a CD44-derived peptide, known to bind the F3b pocket of the FERM domain, starting from the structure with the closed pocket. We compared a simulation without receptor backbone flexibility to a simulation in which receptor backbone minimization was added in the FlexPepDock refinement step of the protocol, to allow for opening of the inaccessible binding site (**Figure 5**). We found that PatchMAN samples near-native fragments on the free FERM domain (with the closed F3b pocket), but cannot identify it as top-scoring on the rigid receptor structure (**Figure 5B**). However, it easily identified the near-native complex structure when backbone minimization was allowed (**Figure 5C,D**). PFPD failed to place the peptide at the closed binding site at the rigid body docking step, requesting an initial opening of the pocket for successful docking (data not shown). This case demonstrates that PatchMAN is able to identify cryptic binding pockets with its sampling approach that takes into account motifs with structural variability. Moreover, PatchMan can open such pockets by positioning the peptide and moving the receptor backbone around it during the subsequent refinement step.

**Figure 5.**
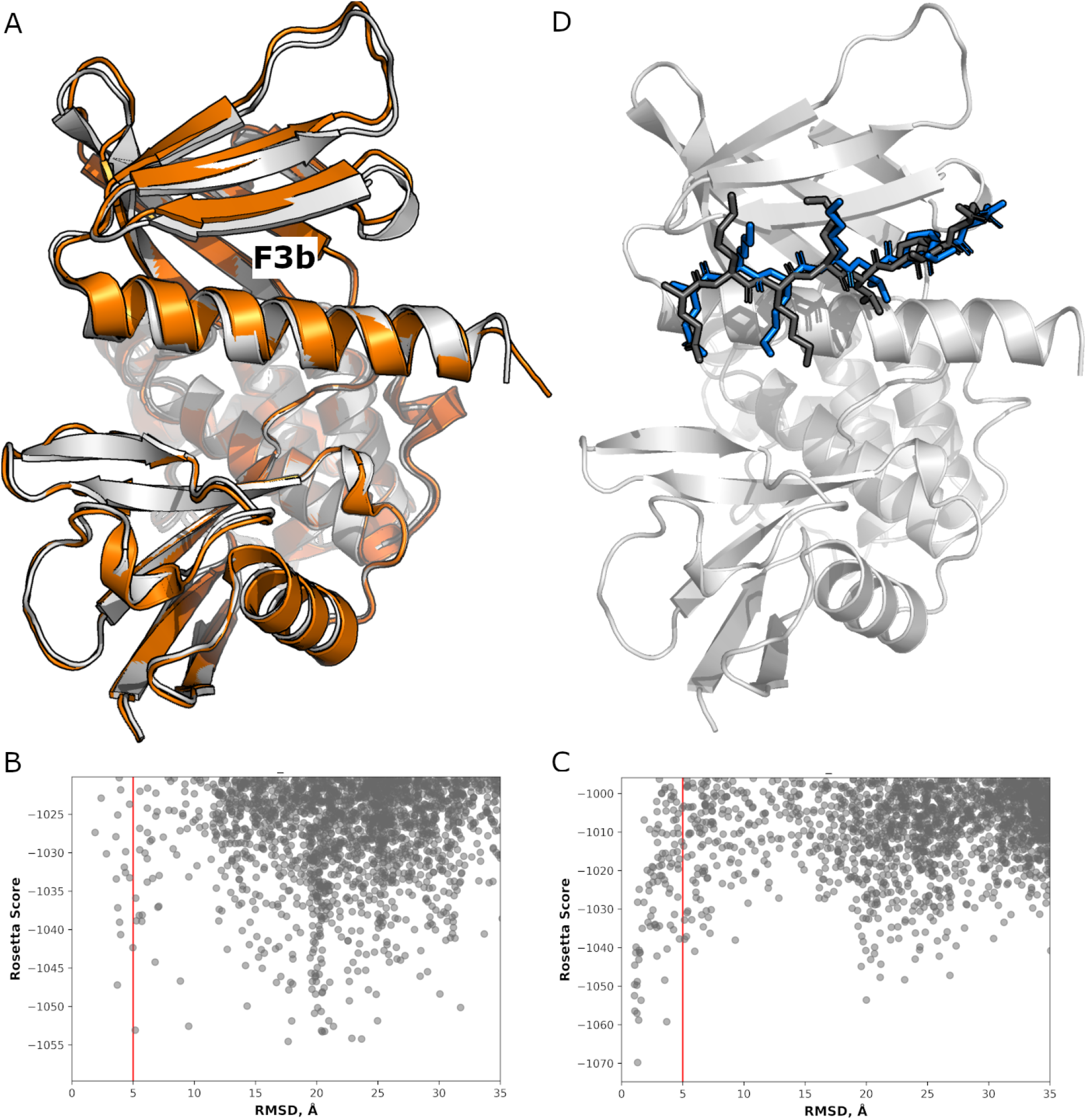
Successful modeling of a peptide into a closed binding pocket using PatchMAN, shown on the example of a CD44-derived peptide binding to the Moesin FERM domain F3b binding site. **A.** The moesin FERM domain structure, showing the unbound closed (grey, PDB ID 1ef1 (29)) and the bound open F3b binding pocket (orange, PDB ID 6txs (30)) structures. A shift in the beta-sheet at the F3b binding site is induced by peptide binding. **B.** PatchMAN simulation without receptor backbone minimization samples the correct binding pocket but misses it at the scoring stage. **C.** PatchMAN simulation including receptor backbone minimization identifies a clear funnel around the native structure. The red line indicates the 5Å RMSD cutoff. **D.** Comparison of model (blue) to crystal structure (grey).

## Discussion

Peptide-protein docking poses particular challenges due to the flexible nature of the peptide partner. The sampling space is vast and complex, as it involves both peptide-internal degrees of freedom as well as rigid body orientation, and often also receptor flexibility (4). Many different approaches were developed to tackle this sampling challenge, usually by breaking it into several smaller, independent sampling steps. However, the biological process of peptide binding is likely to be less modular. It can be seen as a sub-problem of monomer folding, in which the peptide complements the receptor structure, in a way structurally similar to what is observed within other monomers. Here we present a novel, template-based approach that builds upon these biological observations and aims to bridge the gaps between the sampling steps.

PatchMAN leverages information on local structural motifs to search for complementary fragments of the protein surface. These local motifs can be derived from interfaces, but also from completely non-related, monomer structures, as demonstrated in this work. The definition of the structural patches is crucial for success in finding complementing peptides and requires thorough optimization. As was shown previously in *Verschueren et al. (32)*, matching *single* fragments (e.g. using pairs of fragments, one from the receptor and the other from the peptide) instead of *patches* composed of multiple segments (as in PatchMAN) can be useful in some cases, but is not enough to generate a robust sampling strategy. We show that the PatchMAN patch definition is coarse enough to be flexible for conformational changes and thus able to identify near-native templates even from divergent structures, but still specific enough to keep the hit number tractable. We show that the sampling at the binding site is specific to the native-peptide-like conformation, while at the same time diverse, resulting in multiple starting points to local peptide docking, increasing the chances to model a native-like conformation at the refinement step (**Figure 4**).

It is important to note that fast screening of such arbitrary structural motifs is challenging. Here we use MASTER (23): a fast and exhaustive RMSD based search tool that uses the Kabsch algorithm (33) for fast alignment of backbone fragments, managing the growing complexity of multiple segment motif alignments by on-the-fly filtering of non-promising matches. This approach does not include any heuristics, finding all existing alignments within the cutoff RMSD in a matter of seconds, and allowing for fast and efficient sampling.

PatchMAN also demonstrates lower sensitivity to parameters that limit the performance of other methods, such as the peptide length and modeling flanking regions of the peptide. As shown in **Figure 2**, PatchMAN performs equally well on full peptides and on peptide motifs, compared to PFPD performance which decreases when adding the flanking regions. Those findings suggest that using PatchMAN docking can be further improved by connecting templates on the protein surface, to model more complex interactions involving long intrinsically disordered proteins wrapping around a structured partner, a problem only addressed by few studies yet (34).

In PatchMAN, instead of using the sequence of the peptide as the key to modeling its backbone conformation, the focus shifts towards the receptor context. The *receptor* dictates the ensemble of possible peptide structures, making the sampling strategy invariant to peptide sequence. As shown here, most of the selected fragments share very low sequence identity to the docked peptide. PatchMAN opens an avenue for improved peptide design based on these principles. For a targeted receptor pocket, peptide conformations could be extracted and modeled with new sequences. Additionally, peptide backbones could be pieced together to design peptides that interact with multiple adjacent pockets of the receptor.

Moving the focus to the receptor surface also allows for improved modeling by including intrinsic local flexibility (**Figure 5**). For each receptor surface patch, PatchMAN assembles an ensemble of similar motifs from different structures. Hence, even if the surface patch on the receptor is in a “closed” conformation, it can be identified by finding a similar pocket in an “open” conformation. Such pockets can then be opened by superimposing an extracted template followed by short structural refinement. We believe that enriching the hit pool with matches from receptor homologous structures will further improve PatchMAN performance, and specifically may be helpful for cases of conformational changes.

A new era of structural biology has been opened up by Deep Learning, as strongly highlighted by Deepmind’s AlphaFold2 (21). Within this context, we demonstrate in another current study that the peptide docking field can benefit from AlphaFold2 (22), with performance matching our previously developed state-of-the-art PIPER-FlexPepDock protocol (11). The PatchMAN approach presented here performs significantly better, indicating that the local structure information is used more efficiently and accurately than in the current AlphaFold2 version. It remains to be tested how well PatchMAN will perform on structural models of the receptor, and how this work may be optimally incorporated into Deep Learning frameworks.

To summarize, we presented here a robust, quick and high performing global peptide-docking protocol, and demonstrate that the PatchMAN approach is accurate and versatile. As such, it holds high hopes for the peptide modeling as well as peptide design. The incorporation of biological insights and concepts in the development of PatchMAN extends the implications of this work and presents a more general approach to treat peptide-protein docking and binding.

## Methods

### Splitting the surface into structural patches

The protein surface is defined based on solvent accessibility criteria using SASA calculated using the ‘rolling ball’ algorithm implemented in PyRosetta with ball radius = 1.35Å (35). The surface is then split into small structural motifs by selecting the neighbors (Cα-Cα distance within 10Å) around every second surface residue (to reduce the number of overlapping patches). Every motif is defined as one or more disjoint peptide segments, not shorter than 2 amino acids. The maximum length for a single segment is 7 residues for strands and coiled regions, and 11 residues for helices. A stretch is defined to be helical, if it has at least 3 consecutive helical residues based on DSSP (36).

### Searching for local structural motif matches using MASTER

Every patch is searched using the MASTER algorithm (23), against a database of non-redundant protein structures described in the original implementation of MASTER. This database includes 12,661 protein structures (split into X monomers), generated by using BLASTClust (37) at 30% sequence identity on a PDB version of 2014 (23). Briefly, MASTER aligns structural motifs containing multiple disjoint backbone segments to identify all matches within a user-specified RMSD cutoff in a dataset of protein structures. It utilizes the Kabsch algorithm for identifying the match with the lowest RMSD (33), and manages possible combinatorial explosion due to multiple segments in each motif by on-the-fly filtering of partial matches that will not answer the RMSD criteria. For the search, we used the RMSD cutoff of 1.5Å, and took the 50 lowest, as well as the 50 highest RMSD matches to ensure diversity.

### Generating initial complexes for further refinement

For each of the matches we identify the residues that constitute the motif in the corresponding PDB structure. Using PyRosetta (35) we then identify the neighboring peptide stretches (Cα-Cα distance within 8Å), and finally, we elongate peptides longer than 2 amino acids to the desired length in both directions, if possible (**Figure 1B**). Using the rotation-translation (RT) matrices from the MASTER search, the peptide templates are superimposed back onto the receptor protein. We retain those peptides whose backbone does not clash with the receptor (backbone atom distance > 2Å) and who interact with the receptor (at least 45% interacting residues with a heavy atom interaction distance within 5Å). The peptide sequence is then threaded onto the remaining templates (using Rosetta fixed backbone design (38)).

### Refinement of the structures

Rosetta FlexPepDock was used to refine the structures to high resolution and to discriminate near-native models from the rest (as described previously (39)). Here we use the refinement method without receptor backbone minimization for the main benchmark, and refinement with receptor backbone minimization for more challenging targets.

### Criteria for measuring performance

The accuracy of performance was measured as in previous studies (11). In short, the final top 1 percent of the decoys (based on the Rosetta reweighted score (6), using the Rosetta ref2015 scoring function (40)) are clustered and top 10 clusters representatives are analyzed. For the plots in Figure 2 we calculated the number of structures for which the best RMSD model among these 10 representatives lies within the indicated RMSD cutoffs. All results were assessed using RMSD calculated over all interface peptide residue backbone atoms, after superposition of the receptor (i.e., rmsBB_if, as in previous studies, e.g. (11)). Note that the PDB 1LVM was removed from the dataset (since the unbound structure is incorrect), but InterPep2 results are taken from (41) and include this structure.

All scripts and runline commands are provided in the Supplementary Methods section, and on github (https://github.cs.huji.ac.il/alisa/patchman/).

## Acknowledgements

This work was supported, in whole or in part, by the Israel Science Foundation, founded by the Israel Academy of Science and Humanities (grant number 717/2017 to O.S.-F.) and the US-Israel Binational Science Foundation 2015207 (to O.S.-F.). We wish to thank Vasily Khramushin for help in the implementation of PatchMAN.

