## Supplementary material for "PatchMAN docking: Modeling peptide-protein interactions in the context of the receptor surface"

### Supplementary Figures

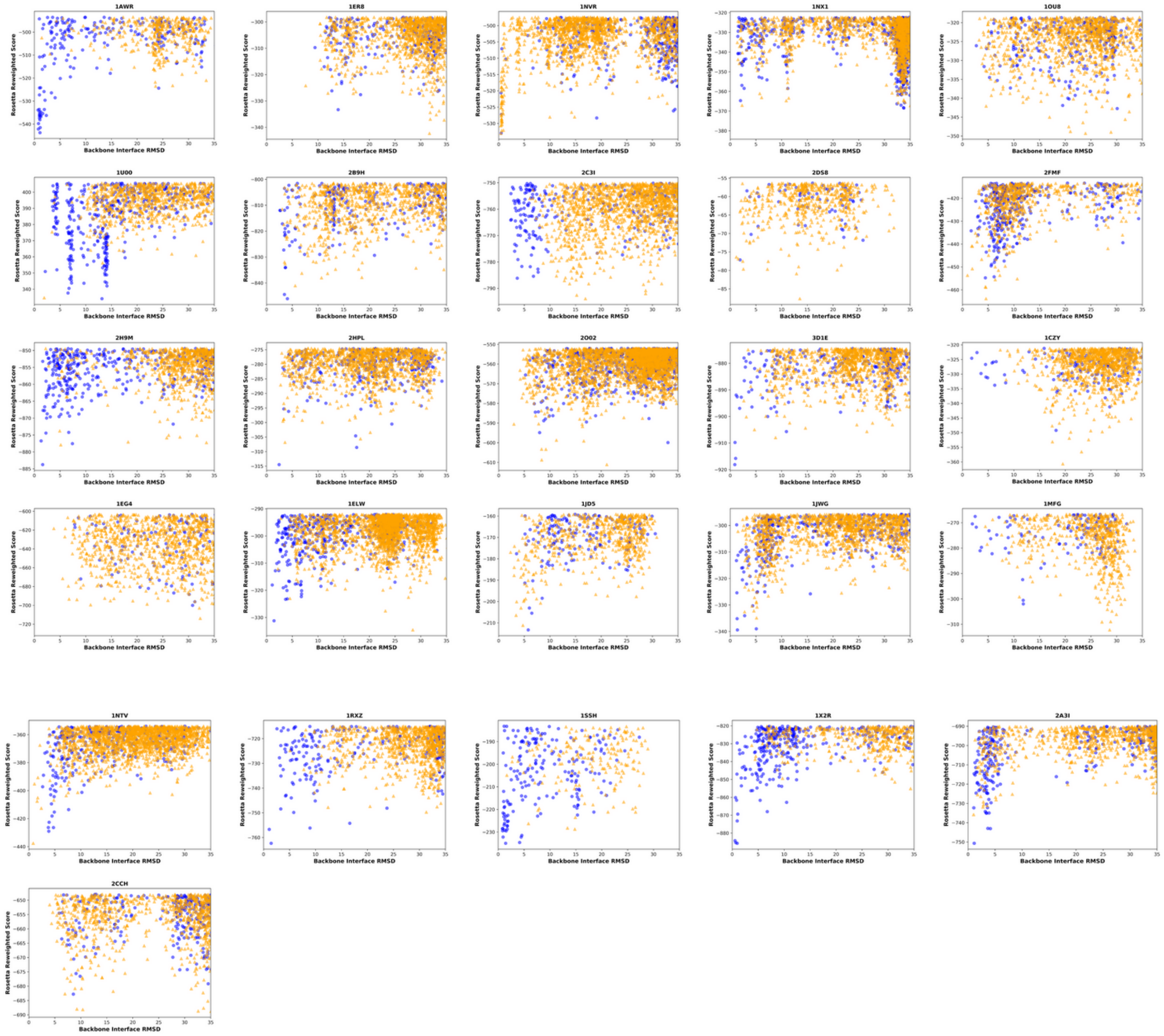


Supplementary Figure S1: Energy landscapes

Rosetta score *vs.* RMSD plots are shown for each of the complexes in the benchmark. Models generated based on templates originating from monomers are indicated in orange triangles, while those stemming from interfaces are indicated by blue circles.

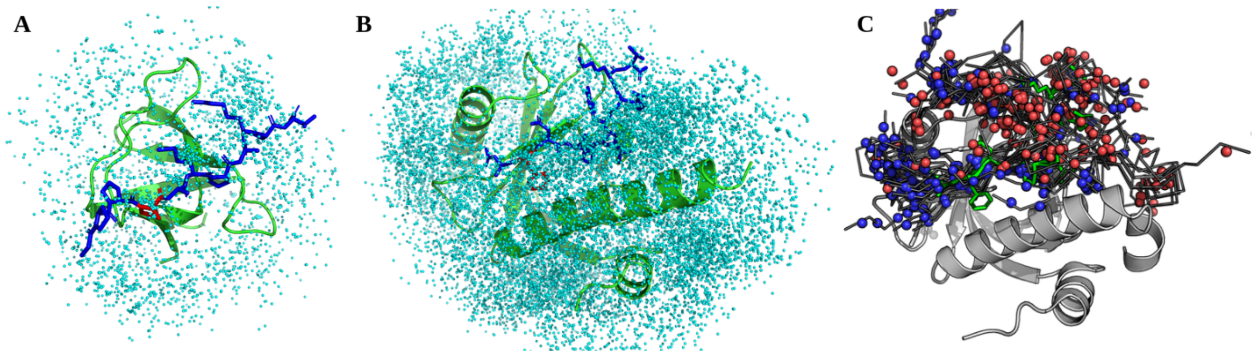


Supplementary Figure 2: Local motif matches map the whole receptor surface

**A-B.** Two examples showing the coverage of the receptor surface with fragments, prior to filtering: (A) 1SSH (42) and (B) 1NTV (26). The receptor and peptide are shown in green cartoon and blue stick representation, respectively. The Cα atoms of one specified peptide residue (highlighted in red in the crystal structure) are shown in cyan spheres. **C.** Sampling at the binding site of 1NTV: Here we show the fragments sampled for a specific surface patch at the binding sites. Fragments are represented by ribbons, with their N and C-termini highlighted with blue and red spheres, respectively. The native peptide structure is shown in green stick representation.
